# Structural Basis for Inhibition of Human Anion Transporter SLC26A6/PAT-1

**DOI:** 10.64898/2026.11.01.757131

**Authors:** Gamma Chi, Tina Bohstedt, Roslin Adamson, Ashley C. W. Pike, Leela Shrestha, Onur Cil, Marc O. Anderson, Maryam Izadiar, Eleanor Williams, Nicola A. Burgess-Brown, Jonathan M. Elkins, Katharina L. Duerr, David B. Sauer

## Abstract

PAT-1, encoded by the gene *SLC2cAc*, is an exchanger of small organic and inorganic anions including oxalate and chloride in the kidney and the intestines. PAT-1 dysfunction is implicated in nephrolithiasis and hyperoxaluria, and its modulation is a potential therapeutic strategy to treat cystic fibrosis. PAT-1 has two small molecule inhibitors: DIDS, a non-specific anion exchanger inhibitor; and PAT1_inh_-B01 which is highly specific to PAT-1. In this study, we report high-resolution cryo-electron microscopy structures of the human PAT-1 homodimer in complex with these inhibitors to reveal their different inhibition mechanisms. These structures also uncover the complex interactions between the transmembrane, STAS and N-terminal domains, with implications on its ion exchange function. Hence, this study provides mechanistic insight into PAT-1 molecular function and inhibition, and contributes to our understanding of SLC26 transport mechanisms.

## Introduction

The SLC26 family are transmembrane proteins that transport small organic and inorganic anions [1], and are involved in a number of physiological pathways and diseases. There are now several experimentally determined structures of SLC26 transporters [2–11], and in each case the protein is composed of N-terminal domain (NTD), transmembrane (TMD) and cytoplasmic Sulphate Transporter and Anti-Sigma factor antagonist (STAS) domains in a domain-swapped homodimer architecture. The TMD has a 7+7 inverted repeat fold, where the central gate domain maintains the dimeric structure while the distal core domain is mobile to facilitate anion transport via an elevator mechanism. The two TMD subunits appear to function independently based on experimental structures of both symmetric (inward and outward) and asymmetric dimers. The substrate-bound SLC26 structures reveal that there are potentially two anion-binding sites [4], termed S1 and S2. Whereas Site S1 is well-characterised via several substrate-bound structures across multiple SLC26 transporters, to date Site S2 has been observed only in SLC26A2 and its function is less clear.

These SLC26 proteins can be inhibited by a variety of chemotypes. These include specific targeting of DRA (SLC26A3) by DRA_inh_-A250 and TQR1122 [12, 13], and nonspecific targeting of the protein by tenidap and UK5099 [14]. Similarly, Prestin (SLC26A5) is inhibited by salicylate [15]. In co-structures of the proteins with each of these compounds [7, 13, 16–22], SLC26 transporters are seen in inward-occluded states. The chemically analogous tenidap and UK5099 appear to act through non-competitively inhibition, whereas DRA_inh_-A250, TQR1122 and salicylate are competitive inhibitors binding directly at the substrate pockets.

Of the SLC26 proteins, Putative Anion Transporter 1 (PAT-1) is encoded by the gene *SLC2cAc* and exchanges various combinations of chloride (Cl^-^), sulphate (SO_4_^2-^), bicarbonate (HCO_3_^-^), formate (HCO_2_^-^), hydroxide (OH^-^) and oxalate (C_2_O_4_^2-^) [23, 24]. The structure of human PAT-1 (PAT-1) was recently determined by cryo-electron microscopy (cryo-EM) with an architecture similar to other SLC26-family transporters [5], in which the TMD mediates ion exchange across the membrane while the STAS domain is proposed to define substrate specificity. The STAS domains interact strongly with large interaction area between them, whereas its two transmembrane domains only limited contacts through their C-terminal TM14 helix, a sign of PAT-1’s reliance on the STAS domain for dimerisation.

PAT-1 has key roles in oxalate and Cl^-^transport in pancreas, intestine and kidney [25–28], and it is a therapeutic target for hyperoxaluria, nephrolithiasis and certain gastrointestinal diseases [29–33]. PAT-1 upregulation in the kidney raises oxalate levels in the renal filtrate [30], suggesting its inhibition is a potential therapeutic strategy to prevent stone formation. There is also evidence that PAT-1 interacts via its STAS domain with other transporters including the chloride transporter CFTR, organic anion transporter NaDC-1, and carbonic anhydrase CA2 [34–36]. Therefore, PAT-1 may be an alternative target to treat cystic fibrosis via indirect modulation or functional circumvention of CFTR.

With its involvement in disease, two PAT-1 inhibitors have been identified for use as research tools and therapeutic proof-of-concept studies, including 4,4-**D**i**I**sothiocyanostilbene-2,2’-**D**i**S**ulphonic acid (DIDS) and PAT1_inh_-B01 [25, 27]. DIDS is a non-specific inhibitor of multiple transporters, competing for the respective sulphate and carboxylate binding sites of the Anion Exchangers (AE) AE1, AE2, and AE3 of the SLC4 family and the SLC13 homolog **I**’m **N**ot

**D**ead **Y**et (INDY) from *Drosophila melanogaster* [37–40]. In contrast, PAT1_inh_-B01 is a highly specific inhibitor of PAT-1 with minimal effect on other SLC26s in both cell-based assays and animal models [25], including the closely related SLC26A9 and DRA (SLC26A3), making it a useful chemical tool for *in vitro* and *in vivo* studies. However, the mechanism of action for PAT1_inh_-B01 remains unknown. Notably, there are no ligand-analogous groups in its three distinctive moieties: **5-ph**enyl (5-Ph) group, tricyclic 2,6,7,11-**t**etraza**t**ricyclo[7.4.0.03,7]**t**rideca-1(9),2,4-trien-8-one (TTT) group, and the 4-**b**romo-5-**m**ethyl-3-(**t**rifluoromethyl)**p**yrazol-1-yl (BMTP) moiety (Figure 1D).

**Figure 1.**
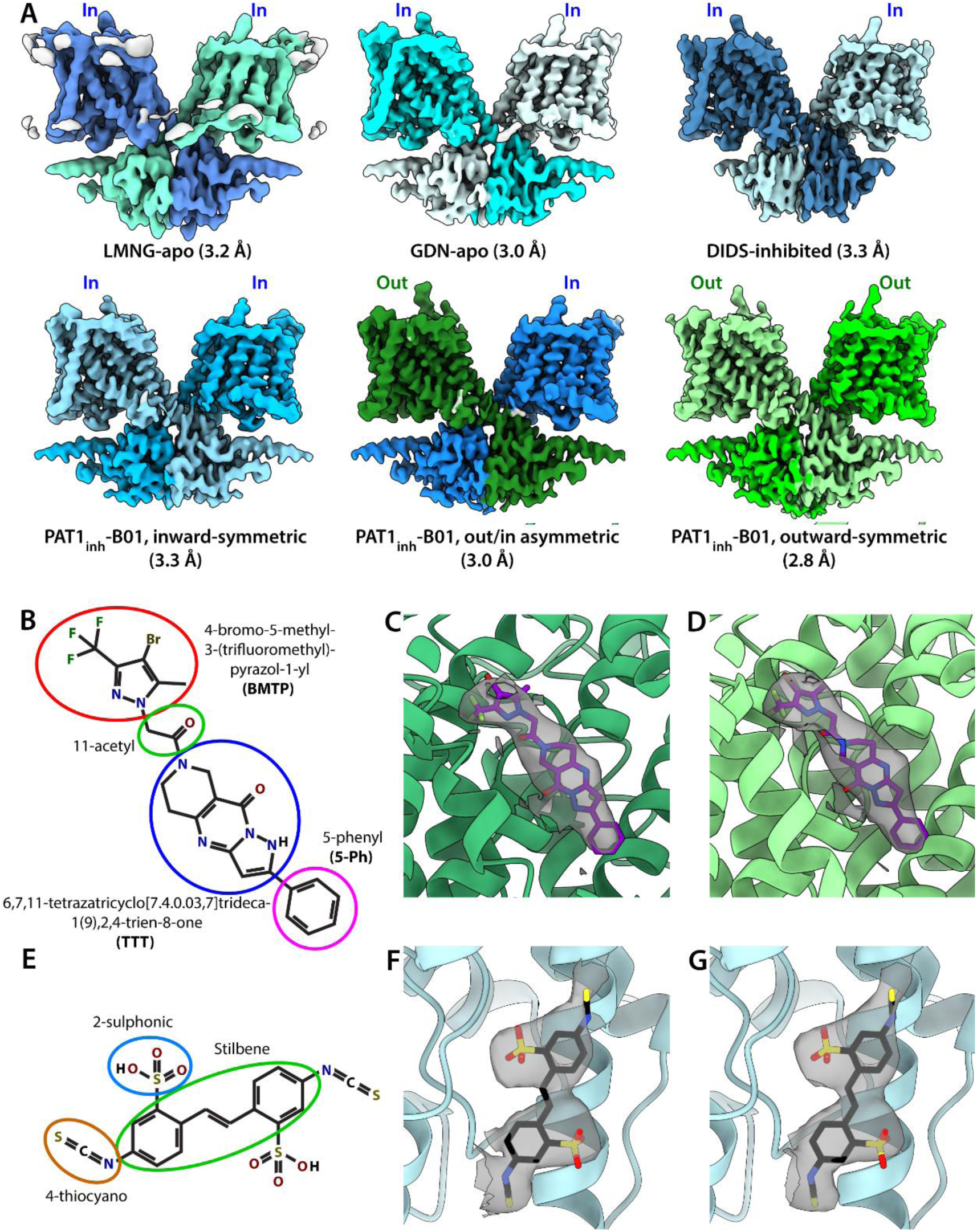
Cryo-EM structures of human PAT-1. **A)** Human PAT-1 (SLC26A6) structures determined in this study. Subunits with inward-open transmembrane domain conformations are represented in blue colours, and those with outward-open transmembrane domains are in green colours. **B)** Structure of inhibitor PAT1_inh_-B01 with defined moieties. **C)** Cryo-EM density of PAT1_inh_-B01 in the outward-open subunit of the asymmetric PAT-1 structure. **D)** Cryo-EM density of PAT1_inh_-B01 in the symmetric outward-open PAT-1 structure. **E)** Structure of DIDS with defined moieties. **F)** Cryo-EM density of DIDS in the primary map. **G)** Cryo-EM density of DIDS in the map where apo-state PAT-1 structure was used as starting reference. The inhibitor density “reappears”, and the model fits into this.

In this work, we present the high-resolution structures of PAT-1 in complex with these two inhibitors determined with cryo-electron microscopy (cryo-EM). The structure of PAT-1 with PAT1_inh_-B01 reveals the molecular basis for the inhibitor’s selectivity toward PAT-1 over its homologues, while the protein’s structure with DIDS structure also adds to the potential of developing SLC26 inhibitors that stabilise SLC26s in inward-open states. Together, these structures will help guide development of therapeutics for nephrolithiasis, as well as other diseases associated with PAT-1 and other SLC26 transporters.

## Results

### PAT1_inh_-B01 stabilises PAT-1 in outward-open state

As a reference for describing the PAT-1 interaction with inhibitors, we first purified and determined the apo-state structure of PAT-1 (Figure 1 A). The full-length human SLC26A6 gene was expressed in HEK293F cells using the BacMam system with a C-terminal Strep-II tag. The cells were solubilized in lauryl maltose neopentyl glycol (LMNG) and cholesteryl hemisuccinate (CHS) and PAT-1 purified using StrepTactin resin. The protein was stable after further purification by size-exclusion chromatography in either LMNG or glycol-diosgenin (GDN), and we therefore determined its structures by Cryo-EM to 3.2 Å in LMNG (LMNG-apo) and 3.0 Å in GDN (GDN-apo). Both PAT-1 structures are symmetric domain-swapped homodimers with both transmembrane domains (TMD) in inward-open (Ci/Ci) states, consistent with the reported PAT-1 structure as well as those of other SLC26s in either apo or substrate-bound conditions [3–5, 8, 9]. Although the buffer for both samples contains 150 mM Cl^-^, the anion-binding Site S1 features a weak density and only in the GDN-apo structure (Supplementary Figure 4 D). As this density was near the noise level of the map, we could not confidently model it as either an anion or a water molecule. This is in contrast to PAT-1, Prestin (SLC26A4) and Pendrin (SLC26A5) structures determined by cryo-EM in similar conditions and conformational states [5–7, 10, 17], where Cl^-^atoms could be resolved at similar map resolutions.

Having resolved an apo state of the protein, we next explored the structural and inhibitory effects of PAT1_inh_-B01 by determining the Cryo-EM structure of PAT-1 in complex with the inhibitor (Figure 1 A). Remarkably, processing of our Cryo-EM dataset revealed multiple conformational states of PAT-1, particularly in its TMD. We separated these in silico during map reconstruction to yield three structures of the PAT-1 homodimer: symmetric outward-open (Co/Co) at 2.8 Å, asymmetric outward-open/inward-open (Co/Ci) state at 3.0 Å, and symmetric inward-open (Ci/Ci) state at 3.2 Å. Of these, the Co/Co state is similar to the previously determined structure of Pendrin in a symmetric outward-open state [6]. The maps were all sufficiently well resolved for model building, and we identified a non-protein density matching the PAT1_inh_-B01 structure in the substrate-binding pocket of subunits in the outward-open conformations (Figure 1 C,D), within both subunits of the Co/Co structure and only the outward-facing subunit of the Co/Ci structure featuring the density. However, such density is missing from TMDs of PAT-1 in an inward-open conformations including the inward-facing subunit of the Co/Ci structure, suggesting that the inhibitor binds only to the outward-open conformation. Given that only the Ci/Ci state is observed for GDN-apo and LMNG-apo structures, the observation of the outward-facing conformation in this dataset suggests that PAT1_inh_-B01 induces conformational change in the protein and stabilises its outward-open conformation.

### PAT1_inh_-B01 selectivity toward PAT-1 against DRA is facilitated by hydrophilic interactions at multiple sites

Within the PAT-1 protein, PAT1_inh_-B01 sits alongside the substrate channel. This sterically blocks substrates from entering the binding site, indicating PAT1_inh_-B01 is a competitive inhibitor of the transporter (Figure 2 E, Supplementary Figure 4 A). Comparison to outward-open anion exchanger AE3 with bicarbonate and DIDS suggests that PAT_inh_-B01’s tricyclic 2,6,7,11-**t**etraza**t**ricyclo[7.4.0.03,7]**t**rideca-1(9),2,4-trien-8-one (TTT) group occupies the likely major substrate-binding Site S1 [38]. Likewise, the inhibitor’s 5-**Ph**enyl (5-Ph) group occupies the deepest groove of the pocket corresponding to the putative minor anion-binding Site S2.

**Figure 2.**
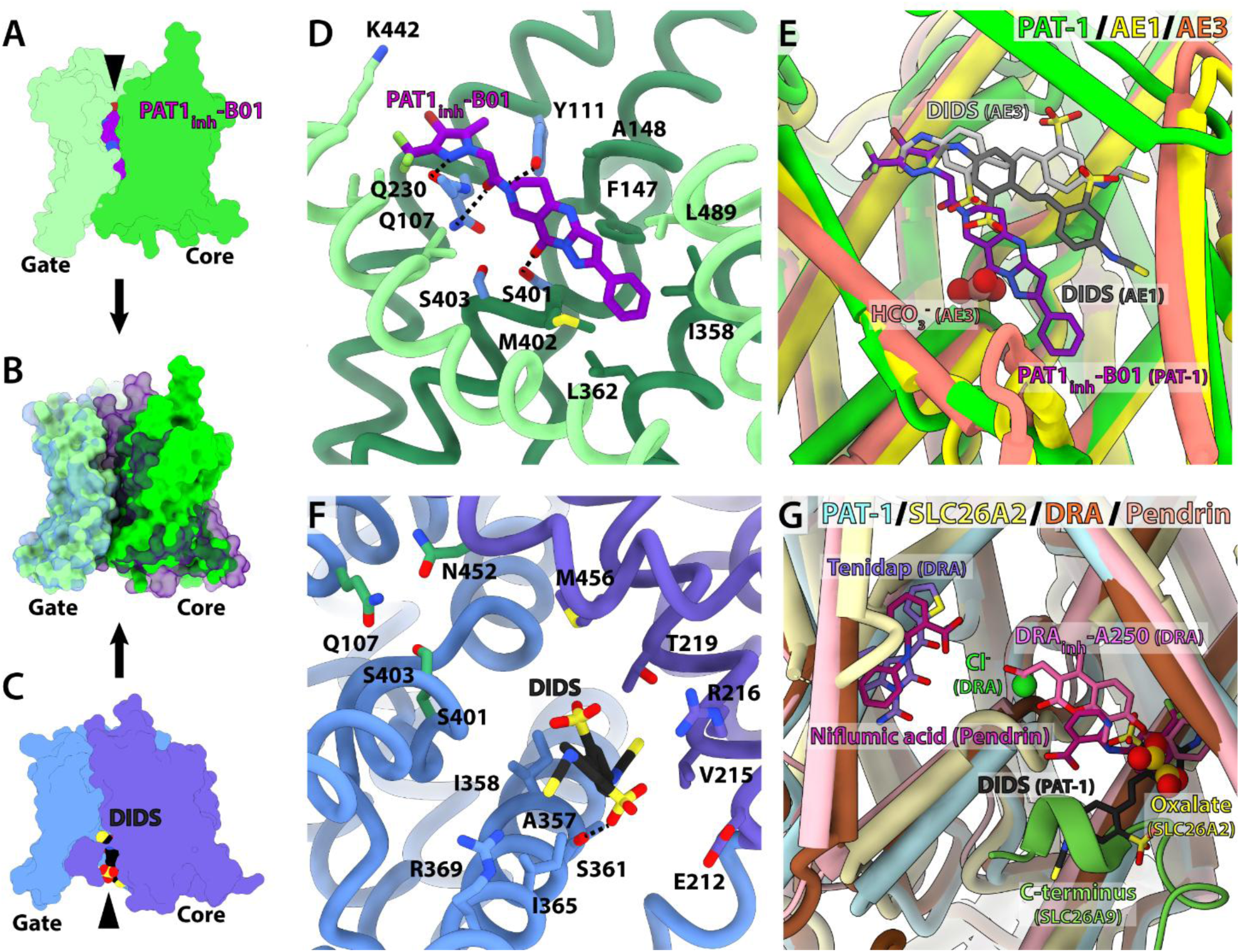
Structures of inhibitor-bound PAT1 transmembrane domain. **A-C)** Surface representations of PAT1_inh_-B01 stabilising PAT-1 TMD in an outward-open conformation (**A**) and DIDS stabilising it in an inward-open conformation (**C**). Overlay of the two structures (**B**) shows rigid-body movement of the core domain whereas the gate domain remains similar between the two states. **D)** PAT1_inh_-B01 binding mode in PAT-1. Light green – gate domain. Dark green – core domain. Light blue – Site S1 residues. The inhibitor makes multiple hydrophilic interactions with Site S1 residues **E)** Alignment of PAT1_inh_-B01-bound PAT-1 structure against DIDS and bicarbonate-bound AE1 (PDB ID: 8T6V) and DIDS-bound AE3 (8Y85), all in outward-open conformations. PAT1_inh_-B01’s TTT group is sited at the equivalent of AE1’s bicarbonate, suggesting that it is occupying substrate binding Site S1. **F)** DIDS binding mode in PAT-1. Light blue – gate domain. Light violet – core domain. Green – Site S1 residues. DIDS is too far from Site S1 to directly replace substrates. **G)** Alignment of DIDS-bound PAT-1 structure against oxalate-bound SLC26A2 (8TNX) tenidap and chloride-bound DRA (8X1S), DRA_inh_-A250-bound DRA (8YRA), and niflumic acid-bound Pendrin (8SHC), all in inward-open conformations. DIDS directly occupies the equivalent of oxalate-binding Site S2 in SLC26A2, but it is too far from Site S1 where chloride is modelled in for SLC26A3.

The PAT1_inh_-B01 binding site has a net positive charge, which would be expected for an anion exchanger substrate pocket. All three of the inhibitor’s moieties are involved in interactions with the protein, suggesting that its high affinity and specificity are not dependent on a single feature. The 5-Ph group makes van der Waals interactions with hydrophobic side chains from I358, L362 and M402 (Figure 2 D). While other SLC26 transporters also have hydrophobic side chains for their equivalent residues, they are still divergent, which likely underlies their differing substrate specificity. The tricyclic ring of the TTT moiety is bounded by Y111 and F147 on one side and V226 on the other. This group has two hydrophilic interactions with the protein: a hydrogen bond between its 8-acetyl group and S401 side chain, and a dipole-dipole interaction between nitrogen on position 6 of the tricyclic ring and the main chain nitrogen of M402. Poor conservation of S401 suggests this residue may partially underlie PAT1_inh_-B01’s specificity within the SLC26 family (Supplementary Figure 5).

The BMTP (4-**b**romo-5-**m**ethyl-3-(**t**rifluoromethyl)**p**yrazol-1-yl) moiety and the 11-acetyl group bridging it to the TTT group have the most complex interactions with the PAT-1 binding pocket residues (Figure 2 D). Of these, the nitrogen atoms from the BMTP moiety and the 11-acetyl bridge make hydrophilic interactions with Q230 on TM5 helix and Q107 on TM1 respectively. Both residues are only partially conserved within the SLC26 family (Supplementary Figure 5), with Pendrin (SLC26A4) being the only member with same amino acids as PAT-1 at equivalent sites, and therefore these amino acids likely contribute to the compound’s transporter selectivity.

### DIDS binds to PAT-1 in inward-open state in the detergent environment

In order to compare the inhibition modes between specific and non-specific inhibitors of PAT-1, we also determined its cryo-EM structure in complex with DIDS (Figure 1 A). The 3.3 Å structure of PAT-1 is in an inward-open conformation, similar to its apo-state structure (RMSD = 0.917 Å for whole proteins, 0.461 Å for TMD).

Within the PAT-1:DIDS complex structure, we found a non-protein density that extend from the substrate-binding pocket to the surface of the TMD (Figure 1 F, Supplementary Figure 4 F). As this region was notably weak, and the overall map at modest resolution, we validated the presence and shape of this density by reconstructing the DIDS-bound PAT-1 map using the GDN-apo map as the starting model. This density reappeared in the final reconstruction, despite it being absent in the starting model (Figure 1 G). This confirmed our identification of this density as corresponding to bound DIDS, though we subsequently used for all analysis the DIDS-bound PAT-1 map generated using the conventional workflow with *ab initio* map generation to avoid any possible model bias.

The portion of this density deepest into the substrate pocket is sufficiently well-defined to allow building an atomic model of the inhibitor. Notably, this site overlaps with Site S2 and binding sites for other SLC26 inhibitors, including DRA_inh_-A250 binding to DRA [18], one of two niflumic acids binding to Pendrin [10], and the self-inhibiting C-terminal peptide of SLC26A9 [8] (Figure 2 G), suggesting this is a hotspot for inhibiting this family.

On the other hand, the other side of this density nearer the surface is poorly defined. Modelling a second DIDS molecule (similar to second niflumic acid molecule in Pendrin) [10], a lipid (similar to tenidap-bound DRA) [20], hexose molecules (present in detergents GDN and LMNG), or a peptide fragment (similar to the C-terminal peptide of SLC26A9) [8] did not yield a convincing fit. Focused 3D classifications suggested sufficient heterogeneity in the density shape to preclude a reliable model building, therefore we opted to leave this region unmodelled.

While the modelled DIDS is surrounded by hydrophilic residues with R216, T219, S361, R369 as well as the backbone amide group between A357 and I358 within 4 Å of the inhibitor, only S361 is close enough for a polar interaction with one of its sulphonic groups (Figure 2 F). Such lack of specific interactions may contribute to the relatively weak density observed in our structure, as well as reports of partial PAT-1 inhibition even at very high DIDS concentrations [41]. The inhibitor’s phenyl stilbene ring is in Site S2, suggesting a competitive mode of inhibition at this site (Supplementary Figure 4 A). Interestingly, the DIDS’ other sulphonic groups, the one not interacting with S361, is oriented toward Site S1 yet 6 – 7 Å away from the anion site. This suggests that, while DIDS is highly effective at competing for Site S2, it may be only somewhat effective at competing with substrates for Site S1.

This PAT-1:DIDS complex map also features weak density in Site S1 (Supplementary Figure 4 E), as previously noted in the GDN-apo map. Though again we have not modelled this density as it is near the noise level of the map, the consistency of this density across multiple maps support the notion it is a water or inorganic anion, and weak or non-competitive binding between DIDS and the species at Site S1.

### PAT-1 structures demonstrate the elevator mechanism of its substrate transport

We next examined the outward-open conformation of PAT1:PAT1_inh_-B01 and the inward-open conformations of PAT1:DIDS and PAT1:apo, noting very little structural change in the protein’s STAS domain and the gate domain of the TMD (RMSD = 0.397 Å and 1.071 Å, respectively). On the other hand, the PAT-1 core domain exhibits a rigid-body translation of 8 – 11 Å across the membrane between the two conformations (RMSD = 0.620 Å). These movements explain the mechanism of action for each inhibitor, where PAT1_inh_-B01 site blocks TM1 and TM3 helices from moving to reach the inward-open state, while DIDS prevents TM8 helix from necessary movements to adopt the outward-open state. Notably, the PAT1_inh_-B01 and DIDS sites both include F147, I358, and M402 despite the inhibitors locking the transporter in different conformations (Figure 2 D,F, Supplementary Figure 5), hinting that these residues are involved in PAT-1’s structural transitions although they do not directly participate in substrate binding.

The rigid-body movement of the core domain within the TMD has different consequences on Sites S1 and S2. PAT-1’s Site S1 is located in a deep groove in inward-open conformations, but nearer the surface in outward-open conformations (Figure 3 B,E, Supplementary Figure 4 A). Whereas N452 and M456 form Site S1 on the gate domain side in inward-open conformations, V226 and Q230 are the gate-side Site S1 residues in the outward-open conformations. This represents a classical elevator mechanism of substrate transport, where the substrate-binding pocket in the core/transport domain “slides” against the gate/scaffold domain (Figure 3 A,D). Therefore, our structures capture snapshots of Site S1-mediated substrate transport by PAT-1.

**Figure 3.**
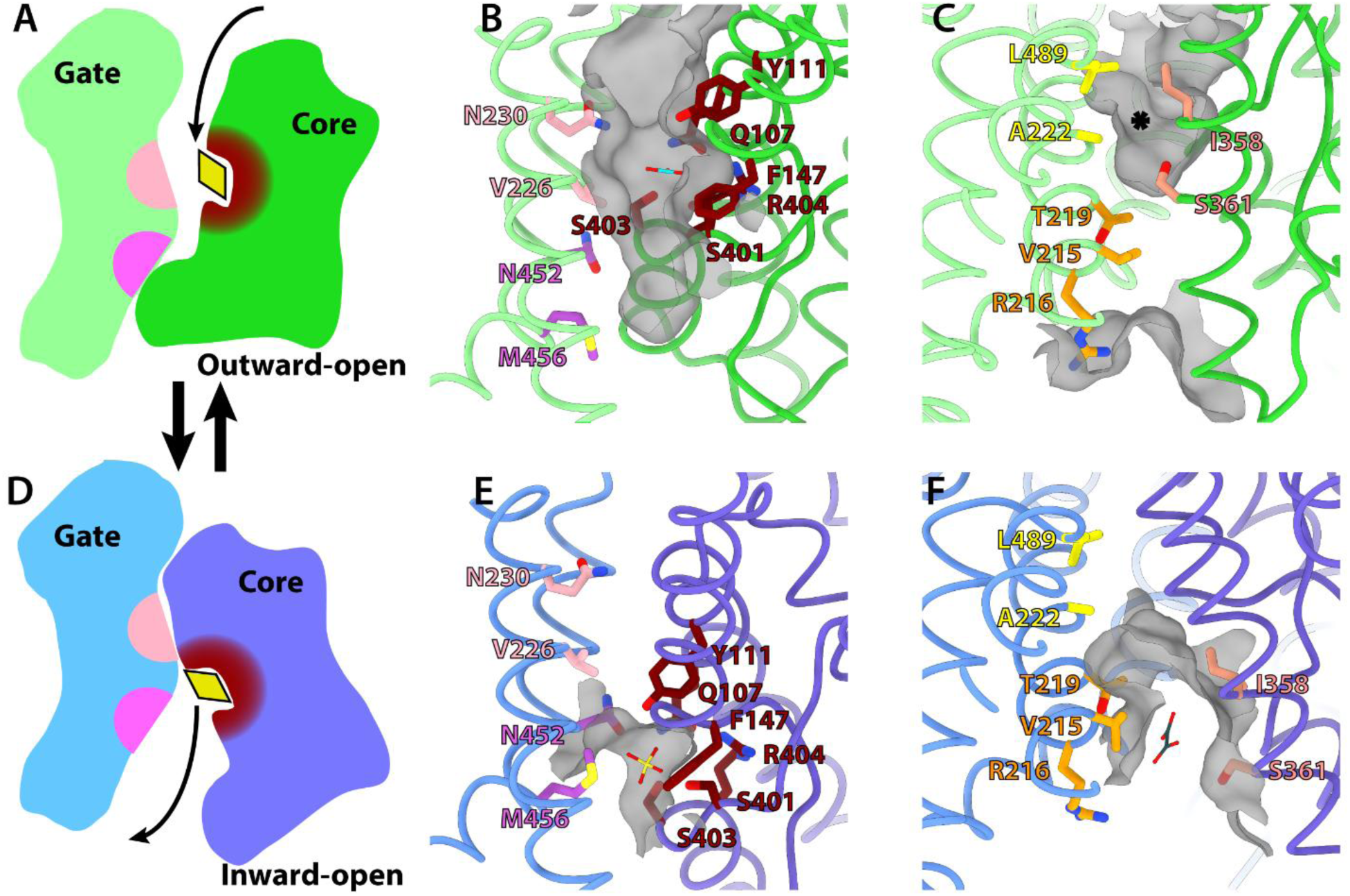
Elevator mechanism of substrate transport by PAT-1. **A,D)** Schematic figure of PAT-1 TMD in outward-open (**A**) and inward-open (**D**) conformations. Surface lining Site S1 on the gate domain side is different between the outward-open (light pink) and inward-open (light violet) states. **B,E)** Structures of PAT-1 Site S1. Dark red – core domain-side Site S1 residues. Light pink – outward-open gate domain-side Site S1 residues. Light violet – inward-open gate domain-side Site S1 residues. Bicarbonate from outward-open AE3 structure (PDB ID: 8Y85) and sulphate from inward-open DRA (8X1T) are overlaid on the PAT-1 structures to denote substrate site. **C,F)** Structures of PAT-1 Site S2. Light orange – core domain-side Site S2 residues. Yellow - outward-open gate domain-side Site S2 residues. Orange – inward-open gate domain-side Site S2 residues. Oxalate from inward-open SLC26A2 (8TNX) and an asterisk are overlaid on the PAT-1 structure to denote substrate site.

On the other hand, the functional role of Site S2 is less clear from our structures. In inward-open conformations, Site S2 is semi-exposed and bounded by V215, R216 and T219 on the gate domain side, and I358 and S361 on the core domain side, analogous to SLC26A2’s oxalate-binding site [4] (Figure 3 F). However, its location in the outward-open PAT-1 is poorly defined. While the core domain Site S2 residues form the deepest groove of the substrate-binding pocket in this conformation, this groove is now bounded by A222 and L489 on the gate domain side (Figure 3 C), making it hydrophobic and S361 alone would not be sufficient for substrate coordination. Hence, whereas Site S2 may be involved in substrate recognition or transport in the inward-open state of PAT-1, its role in the outward-open state may be limited.

### Structural stability of PAT-1’s NTD is correlated with its TMD conformation

One major difference between the various PAT-1 structures is the apparent mobility of its N-terminal domain (NTD), which wraps around the STAS domain near the dimerisation interface (Figure 4 A-F). The cryo-EM density for the NTD is clearest for maps from the PAT-1:PAT1_inh_-B01 dataset, with this region well-defined in Co/Co and Co/Ci maps, and slightly less resolved for the Ci/Ci map. NTD density is only weakly present in the GDN-apo Ci/Ci structure, and it is nearly absent in the PAT-1:DIDS Ci/Ci structure. A 3D refinement using the PAT-1:DIDS particle set and Ci/Ci map leads to a reconstruction with no NTD density, validating this absence as a genuine feature rather than an artefact from starting model or orientation distribution. The apparent presence/absence of the NTD is also unlikely to be a purification artefact; the same purification conditions were used for the GDN-apo and PAT-1:PAT1_inh_-B01 samples until the inhibitor was added immediately before grid preparation, yet the resulting maps have dramatic differences in NTD density. Hence, there is a trend where the NTD’s STAS-bound state is most stable when at least one of two PAT-1 TMDs is in outward-open conformation (i.e. Co/Co and Co/Ci states bound to PAT1_inh_-B01), less stable where inward-open conformations are generally favoured (i.e. Ci/Ci in PAT1_inh_-B01, and apo conditions), and disordered if the TMD is stably locked in an inward-open state by DIDS.

**Figure 4.**
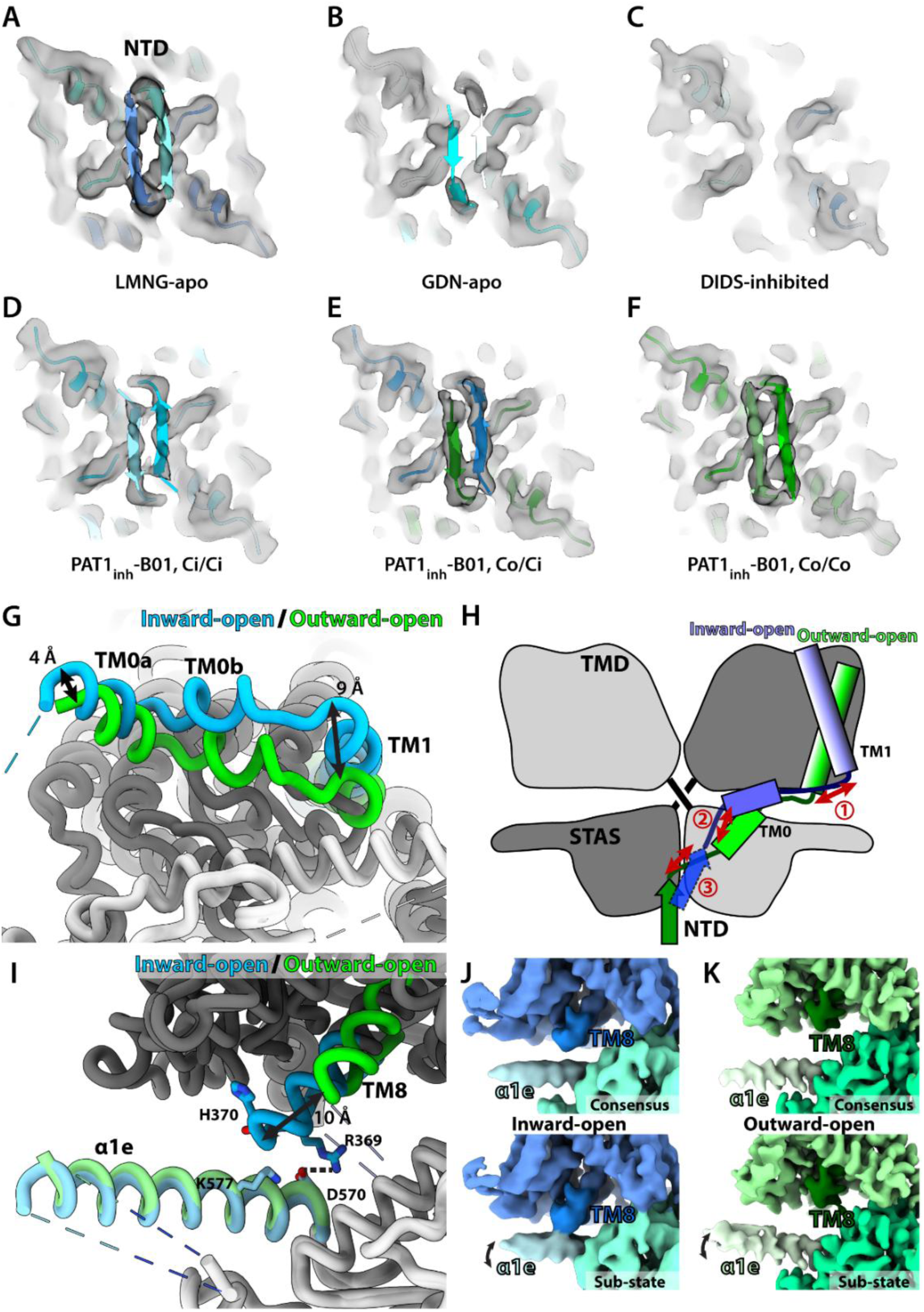
Interactions between PAT-1 TMD and cytoplasmic domains. **A-F)** Cryo-EM densities of PAT-1 NTD in various conditions. Contours were normalised to the levels where detergent belts start to disappear. **G)** NTD-TMD linker shift between inward-open (cyan) and outward-open (green) PAT-1. **H)** Schematic representation of NTD conformational change. ① The 9 Å shift of TM1 helix is correlated with ② 4 Å shift of NTD-TMD linker toward/away from membrane bilayer, and ③ structural instability of NTD. **I)** Interaction between TMD’s TM8 helix and STAS α1e helix. TM8 moves by up to 10 Å toward α1e helix in inward-open conformation, closing their distance to 4 - 6 Å. **J)** Consensus cryo-EM map and sub-state map from 3DVA show that the α1e flexibility brings the two helices further closer to each other in the inward-open state. **K)** Consensus cryo-EM map and 3DVA sub-state map show similar α1e movement for outward-open PAT-1, however this is not enough for interaction with TM8.

This correlation between TMD conformation and the stability of the NTD’s STAS-bound state may be due to the NTD being directly connected to TMD’s core domain (Figure 4 H). The 9 Å shift of TM1 helix between the inward and outward-open conformations leads to a 4 Å move of the NTD-TMD linker away from the NTD (Figure 4 G), exerting tension that would destabilise it. The LMNG-apo structure of PAT-1 indirectly supports this connection, where the TM0 helix position is similar to the outward-open conformation despite its TMD being in an inward-open conformation (Supplementary Figure 4 C). Correspondingly, the NTD is better defined in the LMNG-apo structure than in the GDN-apo and PAT-1:DIDS structures (Figure 4 A-C). Hence, our structures suggest that the NTD-TMD linker mediates the structural relationship between the STAS-bound NTD and the TMD core domain.

### PAT-1 TMD’s TM8 helix interacts with the α1e helix of STAS domain in inward-open conformations

The rigid-body movement of the TMD’s core domain between its inward and outward-open states leads to a ∼10 Å displacement of its TM8 helix (Figure 4 I). As a result, the distance between TMD TM8 helix and STAS α1e helix is ∼4 Å in inward-open conformations but ∼15 Å in outward-open conformations. We therefore assessed whether there are at least transient but direct interactions between the TM8 and α1e helices by analysing Co/Co and LMNG-apo PAT-1 structures with 3D variability analyses on the STAS domains. In some modes, the α1e helix shows mild rigid-body variability of up to 3 Å, rotating with its N-terminus as the pivot point in both Co/Co and LMNG-apo particle sets, while the rest of STAS domain is largely static in all modes. Whereas such positional variability is not enough for TM8/α1e interactions in the Co/Co structure (Figure 4 K), this is sufficient for direct interactions between the two helices in the inward-open LMNG-apo structure (Figure 4 J).

## Discussion

### SLC26 transporters have diverse modes of inhibition

Our PAT-1 structures with PAT1_inh_-B01 and DIDS reveal how this protein is inhibited by two chemotypes, and add to the growing number of inhibitor-bound SLC26 structures. The PAT-1:PAT1_inh_-B01 complex structure is the first reported instance of an SLC26 protein being stabilised in an outward-open conformation by an inhibitor. An inhibitor driving the target transporter into a non-resting conformation has also been observed for cation-chloride co-transporters (SLC12s) [42, 43], anion exchangers (SLC4s) [38–40], and divalent anion/sodium symporters (SLC13s) [37]. With SLC26-family transporters, however, inhibitor-bound structure reported to date have typically been stabilised in inward-facing or occluded states [7, 13, 16–22]. Therefore, our structure of PAT-1 in complex with PAT1_inh_-B01 and DIDS demonstrate that SLC26 family transporters can be inhibited in multiple conformations.

Interestingly, DIDS binds on the cytoplasmic side without inducing a conformational change on PAT-1. This was unexpected, as DIDS binds to the extracellular side of outward-open proteins in all other structures of the molecule with anion transporters [37–40] (Figure 2 E). As seen with the PAT1_inh_-B01 complex structure, PAT-1 in GDN detergent supports sufficient structural flexibility for PAT-1 to change its conformation and accommodate ligands. Therefore, detergent-induced structural rigidity is unlikely to force DIDS binding to only the inward-facing conformation. This raises two possibilities: 1) this structure is representative of physiological inhibition, where DIDS stabilises PAT-1 in the inward-open conformation, reaching the cytoplasm through an unknown mechanism; or 2) the PAT-1:DIDS complex structure is an artefact caused by detergent solubilisation and the physiologically relevant binding site for DIDS is on the extracellular side of the protein. The first hypothesis is consistent with other SLC26 inhibitors, most of which also bind from the cytoplasmic side (except PAT1_inh_-B01) [7, 13, 16–22]. However, this requires the membrane-impermeable DIDS to cross the plasma membrane via an as-yet unknown transport mechanism. The second hypothesis is consistent with DIDS’ inhibition mode for other anion transporters [37–40], but this does not explain the lack of an outward-open subpopulation in the cryo-EM dataset.

Although PAT-1:DIDS is in an inward-open state in our structure, the DIDS binding mode is in line with its common inhibition mechanism for anion transporters in general. As reported with DIDS-bound AE3 and INDY structures [37–40], the molecule binds to the substrate-binding pocket with its sulphonate group 4 – 6 Å from their canonical anion-binding site functionally equivalent to Site S1 in PAT-1. In the case of AE3, DIDS binding does not preclude substrate binding (Supplementary Figure 4 A), as a bicarbonate molecule is observed in Site S1. Likewise, we observe a weak density at Site S1 of our PAT-1:DIDS structure (Supplementary Figure 4 E). This further supports the possibility that DIDS is a non-competitive inhibitor for multiple anion exchanger families.

### Accessory sites in the transmembrane and cytoplasmic domains may contribute to anion transport by SLC26 transporters

Our PAT-1 structures add to the growing evidence of Site S2’s involvement in ion transport by SLC26 proteins. The role of Site S1 in substrate binding is well-characterised for anion transporters of SLC4, SLC13 and SLC26 families featuring elevator mechanism of substrate transport [3, 6, 39, 40, 44, 45], where it facilitates import/export of anions across the cell membrane. On the other hand, the structure-function of Site S2 is less-understood. Site S2 of SLC26A2 is capable of binding substrate anions such as oxalate [4], and our PAT-1:DIDS structure adds to DRA:DRA_inh_-A250 and Pendrin:niflumic acid structures as Site S2-occupying inhibitors [10, 18] (Figure 2 G, Supplementary Figure 4 A), suggesting this is a site of competitive inhibition. This raises two possible molecular models for anion exchange by SLC26 transporters: 1) substrate and conformation-dependent binding model as proposed for SLC26A2; and 2) the central/accessory sites model where Site S2 attracts anions for faster substrate binding or release at Site S1, similar to the model proposed for AE1. The first model is supported by the SLC26A2:oxalate structure and the observed ability of Site S1-compromising PAT-1^R404V^ mutant to retain chloride import [4, 5]. The second model is consistent with most experimental structures of anion exchangers featuring substrates only at Site S1, regardless of substrate type or conformational states. Our structures appear to support the second model. Whereas Site S2 in inward-open conformation is lined by hydrophilic residues capable of binding anions, this site is likely too hydrophobic for substrate binding in the outward-open conformation. Hence, we speculate that Site S2 has an accessory role in substrate export, for example attracting anions in the inward-open state, but that it has very limited or no role in import. Site S1 would primarily facilitate both import and export in this model. Similar differences in the existence and functional importance of a second binding site have been observed for the LeuT family, where the significance of Site S2 is unclear and may vary between homologues [46–49].

Our cryo-EM structures also highlight the mobility and potential functional role of PAT-1’s NTD. The NTD is clearly bound to the STAS domain in the outward-open PAT1_inh_-B01-bound state. Its density is however much weaker in inward-open state structures, suggesting that its mobility is correlated with the TMD conformation. Flexibility in the NTD appears to be mediated by the NTD-TMD linker that leads to the core domain, whose S1 helix shifts as much as 9 Å between inward and outward-open conformations. This steric “tug-of-war” between the NTD and TMD also appears to be a reversible feature, since the PAT-1:PAT1_inh_-B01 sample went through the apo state during purification where the NTD would have been mobile, yet the later addition of the inhibitor leads to the NTD clearly bound to the STAS domain.

Interestingly, similar NTD-disordered states have been reported for other SLC26 transporter structures. All structures of wild-type human and dolphin Prestins (SLC26A5) feature an ordered NTD bound at the STAS dimerisation interface [16, 17]. However, the NTD is not resolved in thermostabilised human Prestin structures regardless of their conformation [7]. Likewise, NTD density is missing in SLC26A7 and SLC26A11 structures [3, 11]. Such reversible NTD binding has been reported in other SLCs, including cation-chloride co-transporters (CCCs) where the NTD interacts with either the TMD to auto-inhibit transport function or the cytoplasmic C-terminal domain (CTD) [50, 51]. The functional importance of the NTD in some SLC26-family transporters is demonstrated by pathogenic mutations in this region of Pendrin that lead to Pendred Syndrome [52, 53]. While its molecular function in the transport cycle and regulation of the SLC26 transporters, and PAT-1 in particular, remains to be elucidated, our results add to the growing evidence for a potential regulatory role of cytoplasmic NTD and STAS domains in SLC26 function.

Our structures also reveal the extensive network of interactions between the TMD and STAS domains of PAT-1, and of SLC26s in general. PAT-1 features two domain swaps: one between NTD and TMD, then another between TMD and STAS. There are also conformation-dependent TM8-α1e helix interactions, and stable interaction clusters near the symmetry axis of the protein (Supplementary Figure 4 B). These interaction points for PAT-1 are also reported for other SLC26 transporters [8, 10], and likely contribute to the general stability of their TMD/STAS domain architecture. While many dimeric SLCs feature structured soluble domains, they are usually inherently flexible relative to the transmembrane domains (SLC4, SLC9A) [38–40, 44, 54, 55] or undergo significant conformational changes between structures (SLC12, SLC53) [42, 43, 50, 51, 56–58]. In this sense, SLC26s are unique for the stable inter-domain structure with NTD being the only mobile feature. While requiring further study, one functional role of such stability may be in substrate binding and specificity as proposed with the crystal structures of Prestin STAS domain [59].

Overall, our structural studies of PAT-1 make important contributions to our understanding of SLC26 molecular function and inhibition while also raising new questions. These will form the basis for new molecular studies as well as SLC26-targeting therapeutics development for diseases associated with PAT-1 and beyond.

## Methods

### Synthesis of PAT1_inh_-B01

PAT1_inh_-B01 has been reported previously as a hit molecule from high throughput screening [25]. Herein we report the synthesis of this molecule (Supplementary Figure 7, Supplementary Methods). The synthesis begins with acid-catalyzed condensation between 3-amino-5-phenylpyrazole (**1**) and 1-*tert*-butyl 3-methyl 4-oxopiperidine-1,3-dicarboxylate (**2**), promoted by acetic acid, to generate Boc-protected intermediate **3**. Next, acid-catalyzed deprotection of **3** was accomplished using HCl (1M in EtOAc) to generate the amine product (**4**) conveniently as a hydrochloride salt. Finally, this molecule was couple under EDCI-mediated conditions to 4-bromo-5-methyl-3-trifluoromethyl-pyrazol-1-yl)-acetic acid (**5**) to generate the product. Notably, none of the steps required chromatographic purification, and the final product could be isolated in ∼95% purity by a simple trituration step.

### Protein Expression, Purification and Cryo-EM Sample Preparation

Codon-optimised gene for isoform 1 of human SLC26A6 (which codes for PAT-1) was ordered from Genscript (USA) and cloned into pHTBV1.1 vector with C-terminal 3C protease site, green fluorescent protein (GFP) tag and Strep-II tag. *Escherichia coli* cells (DH10Bac) were transformed with this plasmid, from which bacmids were extracted. Sf9 cells (ATCC, USA) grown in Sf9-II media (Gibco, USA) were transfected with the bacmid to generate BacMam baculovirus carrying the recombinant SLC26A6 gene. The baculovirus stock was added to HEK293-F “Expi” cells (Thermo Fisher Scientific, USA) at 2.0 x 10^6^ cells/mL at 3 % v/v titre, then the culture was grown at 37°C, 8 % CO2 and 80 % humidity for 40 hours before harvest with a centrifuge set to 900*g* and 14°C for 10 minutes. The cell pellets were flash-frozen with liquid nitrogen then stored at -80°C until purification.

For purification, the cell pellet was thawed, then resuspended in buffer containing 50 mM 4-(2-hydroxyethyl)-1-piperazineethanesulfonic acid (HEPES) pH 7.5, 150 mM NaCl, 1 % LMNG and 0.1 % CHS for solubilisation. After 1 hour incubation, the slurry was centrifuged at 45,000*g* and 4°C for 90 minutes to remove debris. The supernatant was incubated with StrepTactin-SF resin for 2 hours at 4°C, then the resin was collected with a gravity flow column. The resin was washed three times with 5 column volume (CV) of purification buffer A (20 mM HEPES pH 7.5, 150 mM NaCl, 0.005 % LMNG, 0.0005 % CHS). Protein was eluted from the resin with 3 CV of the purification buffer A supplemented with 5 mM D-desthiobiotin. The purification tag was cleaved from the PAT-1 protein by incubation with 3C protease overnight at 4°C.

For the LMNG-apo sample, the affinity-purified PAT-1 was further purified by size-exclusion chromatography on a Superose 6 10/300 Increase column (Cytiva, USA) equilibrated with purification buffer A. For the GDN samples, 5 % GDN stock was added to the affinity-purified PAT-1 to a final concentration of 0.1 %, then it was subjected to size-exclusion chromatography with a column equilibrated with purification buffer B (20 mM HEPES pH 7.5, 150 mM NaCl, 0.02 % GDN).

To determine the structures of PAT-1 in the presence of inhibitors, DIDS or PAT1_inh_-B01 in DMSO were added to SEC-purified protein in GDN to a final concentration of 400 μM and 250 μM, respectively. All protein samples were concentrated to ∼ 3 mg/mL (18 µM), then they were applied to Quantifoil (SPT Labtech, UK) 300-mesh gold grids 1.2/1.3 hole-spacing with Vitrobot Mark IV (Thermo Fisher Scientific, USA).

### Cryo-EM Data Collection, Processing and Model Building

All cryo-EM grids were clipped with autogrid ring and c-clips (Thermo Fisher Scientific). They were assessed for quality with Glacios equipped with Falcon 3 detector (Thermo Fisher Scientific) at Oxford Particle Imaging Centre (OPIC), University of Oxford, United Kingdom. High-resolution cryo-EM datasets were collected, including some micrographs collected with 30° stage tilt for the LMNG-apo and PAT-1:PAT1_inh_-B01 datasets to overcome preferred orientations, at either OPIC or electron Bio-Imaging Centre (eBIC), Harwell, United Kingdom. Details of collection statistics are available on Supplementary Table 1.

All cryo-EM datasets were processed and maps calculated using standard single particle reconstruction workflows in various versions of Cryosparc^TM^ (Structura Biotechnology, Canada) and Relion v5.0 (Supplementary Figures 1, 2 and 3) [60, 61].

For the PAT-1:DIDS dataset, final particles from Cryosparc were exported to Relion for Blush Refinement to realign them and reconstruct a map with less artefact from preferred orientations, then they were returned to Cryosparc for a local-refined with a mask covering the full protein to further align them. The particles were also refined with the Ci/Ci structure from PAT-1:PAT1_inh_-B01 dataset, which has the least preferred orientation artefact of all inward-open structures, to validate the presence of DIDS density as well as the absence of NTD density.

The PAT-1 structure model in the LMNG-apo condition was generated by first creating an initial model by replacing equivalent residues of SLC26A9 (PDB ID: 6RTC) [9] with Chainsaw on CCP4i [62]. It was manually built and refined with Coot [63], then auto-refined with Phenix [64]. GDN-apo, PAT-1:DIDS, and Ci/Ci PAT-1:PAT1_inh_-B01 structure models used this LMNG-apo structure as a starting model, with further manual reconstruction in Coot and refinement in Phenix. Ligands and restraints were generated with AceDRG on CCP4i [65]. The outward-open PAT-1 structure model was generated by replacing equivalent residues of outward-open Pendrin (7WLE) [6] with Chainsaw on CCP4i. This model was then manually rebuilt with Coot, then auto-refined with Phenix.

Structure figures were generated with UCSF ChimeraX v1.7 and v1.8 [66].

### Sequence Alignment

Protein sequences for the canonical isoforms of human SLC26 transporters were downloaded from Uniprot. The sequences were aligned with Clustal Omega [67], then were visualised with ESPript 3 [68].

## Data Availability

All structure maps and models are available on Protein Data Bank (PDB) and Electron Microscopy Data Resource (EMDR). These are: apo-state PAT-1 in LMNG (PDB ID: 32XU, EMDR: 59257), apo-state PAT-1 in GDN (32XQ, 59253), Ci/Ci PAT-1:PAT1_inh_-B01 (32XS, 59255), Ci/Co PAT-1:PAT1_inh_-B01 (32XT, 59256), Co/Co PAT-1:PAT1_inh_-B01 (32XR, 59254), and PAT-1:DIDS (32XV, 59258).

## Acknowledgement

We thank Elizabeth Maclean, Dr. Loic Carrique, Dr. Helen Duyvesteyn and Dr. Thomas Walters at Oxford Particle Imaging Centre (Oxford, UK) for their assistance with electron microscopes. We also acknowledge Diamond for access and support of the cryo-EM facilities at the UK national electron Bio-Imaging Centre (eBIC), proposal BI41204. We also thank Dr. Brian Marsden at Centre for Medicines Discovery (Oxford, UK) for his assistance with cluster maintenance. This research was carried out with funding from the Innovative Medicines Initiative 2 Joint Undertaking (JU) under grants agreement No. 875510 (EUbOPEN). The JU receives support from the European Union’s Horizon 2020 research and innovation programme and EFPIA and Ontario Institute for Cancer Research, Royal Institution for the Advancement of Learning, McGill University, Kungliga Tekniska Hoegskolan, and Diamond Light Source Ltd. This research was also funded in part by the Wellcome Trust (Grant No. 106169/Z/14/Z), and grants from NIH (DK126070, EY036139) and Cystic Fibrosis Foundation. Oxford Particle Imaging Centre was funded by a Wellcome Trust JIF award (Grant No.060208/Z/00/Z) and is supported by equipment grants from WT (093305/Z/10/Z).

## Author Contributions

D.B.S., K.L.D, J.E. and G.C. designed this project. T.B. and L.S. performed molecular biology, with N.A.B.B., K.L.D, J.E. and E.W. supervising them. G.C, T.B. and R.A. performed protein expression. G.C., T.B. and R.A. performed protein purification. M.O.A., M.I and O.C. performed PAT1_inh_-B01 synthetic chemistry and chemical characterisation tasks. G.C., T.B. and R.A. carried out cryo-EM sample preparation, data collection and data processing. G.C. refined structure models, which were proofread by A.C.W.P. for internal validation. G.C. prepared the manuscript. G.C. and D.B.S. revised the manuscript.

## Supplementary Information

**Supplementary Table 1.**
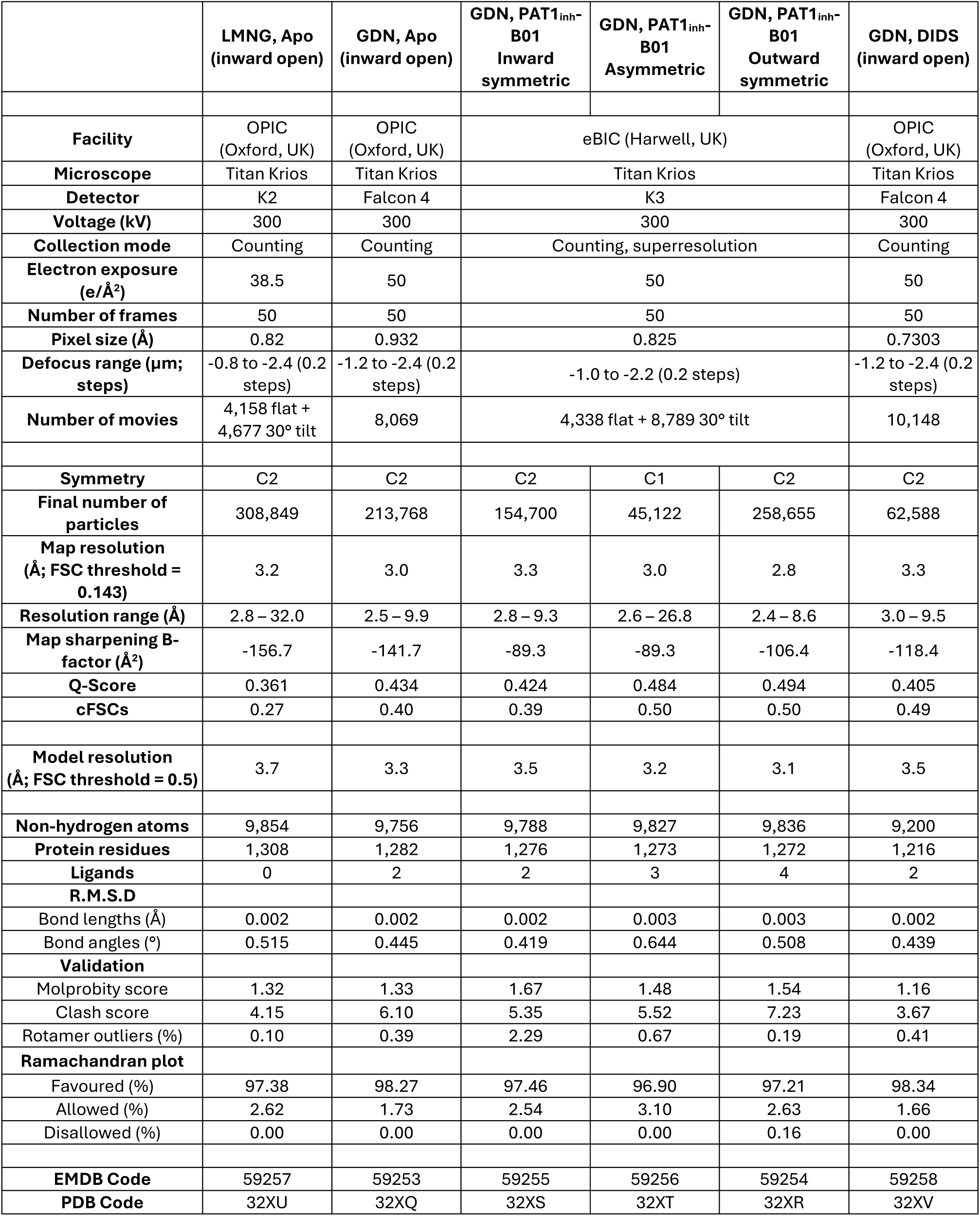
Data table of presented PAT-1 cryo-EM structures.

**Supplementary Figure 1.**
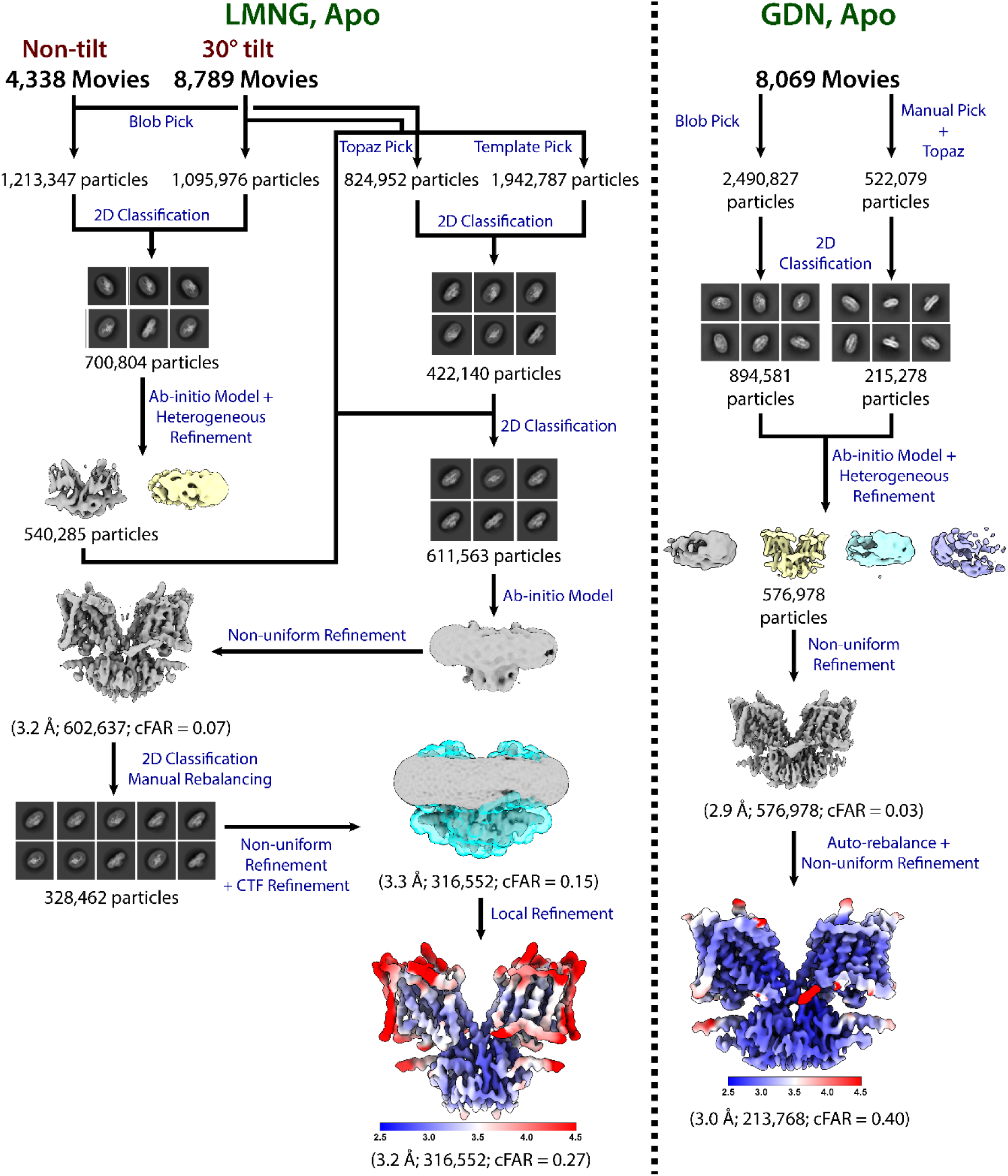
Cryo-EM data processing workflow for LMNG-apo and GDN-apo PAT-1.

**Supplementary Figure 2.**
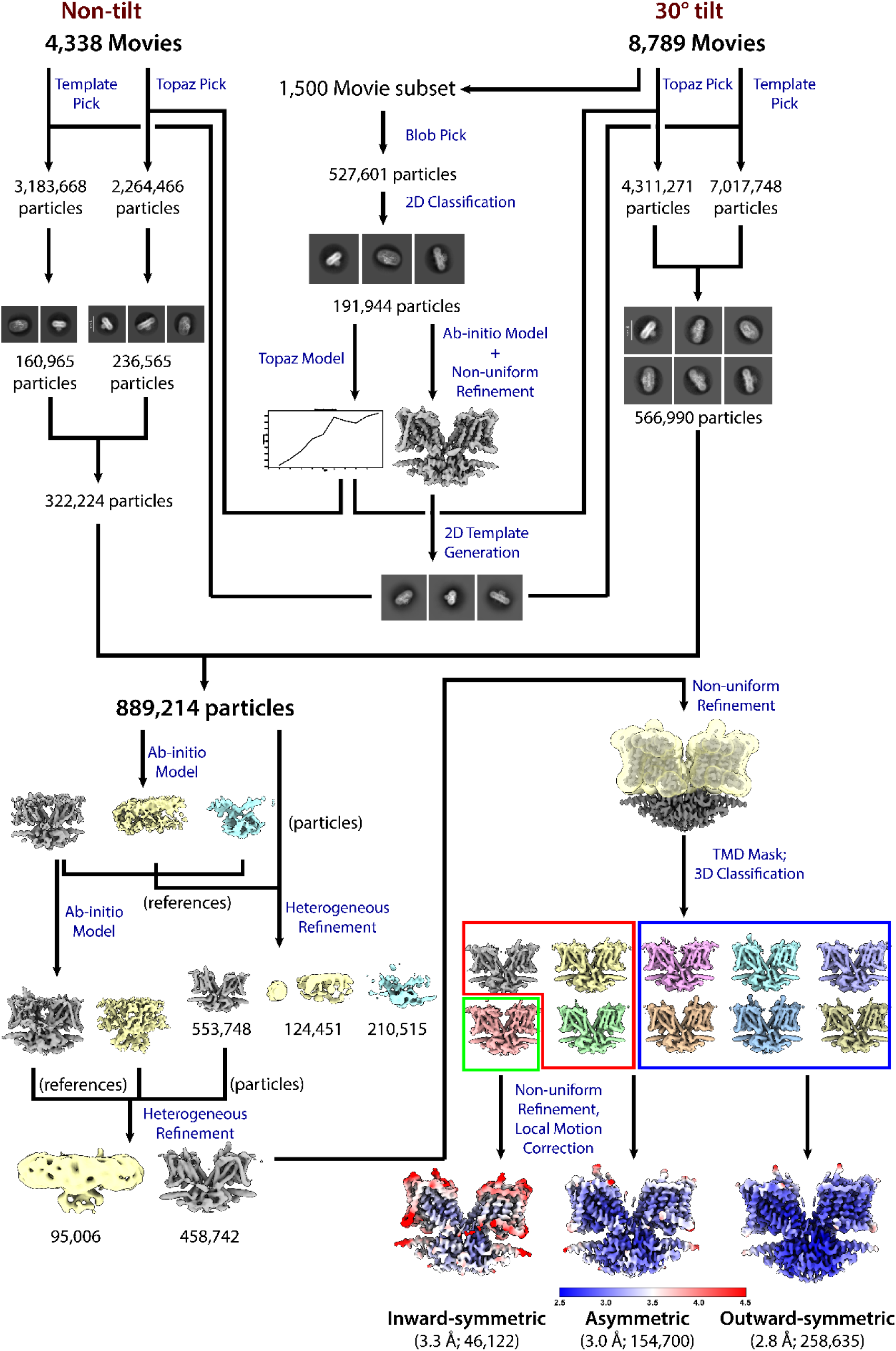
Cryo-EM data processing workflow for PAT1_inh_-B01-bound PAT-1.

**Supplementary Figure 3.**
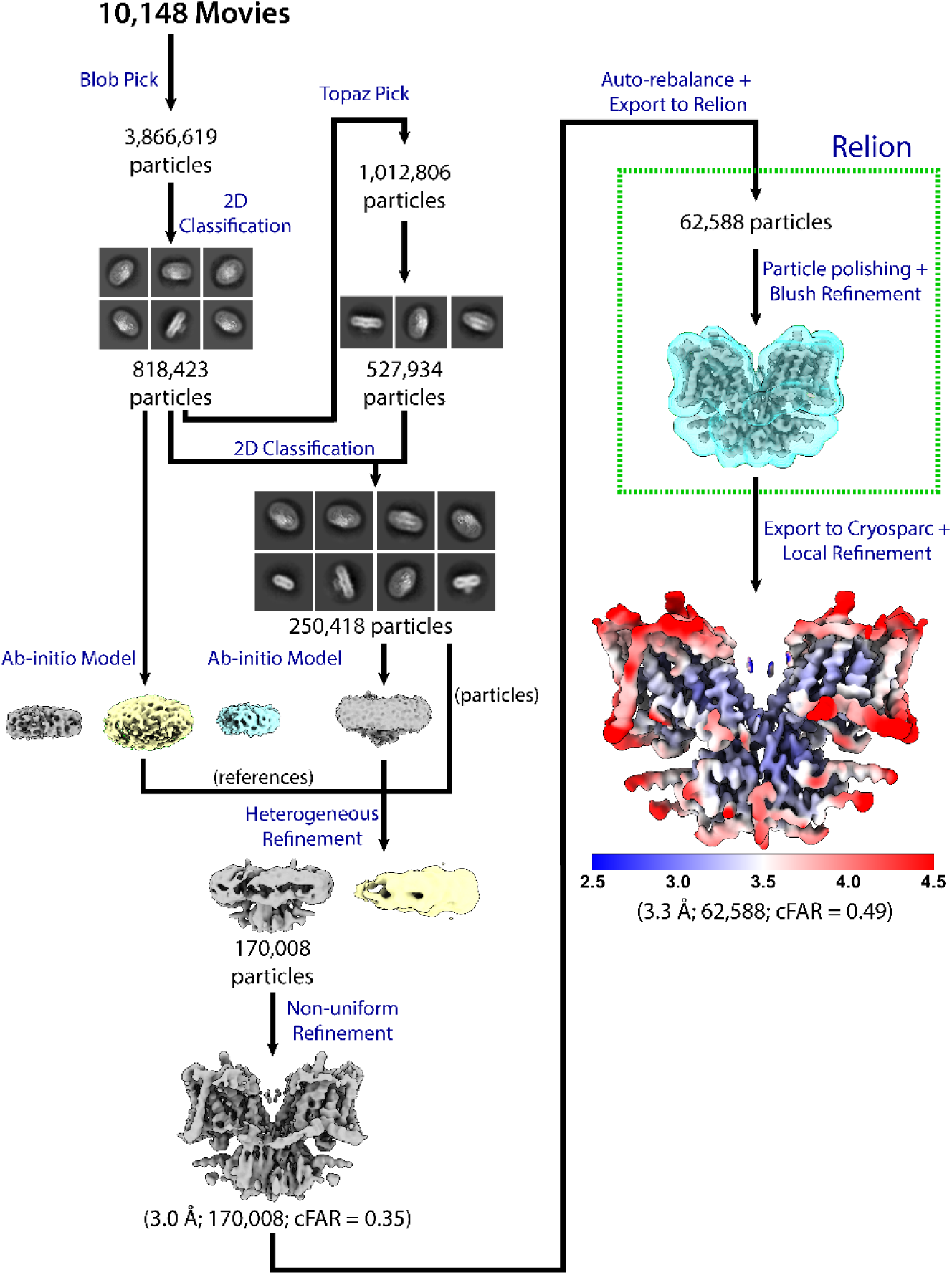
Cryo-EM data processing workflow for DIDS-bound PAT-1.

**Supplementary Figure 4.**
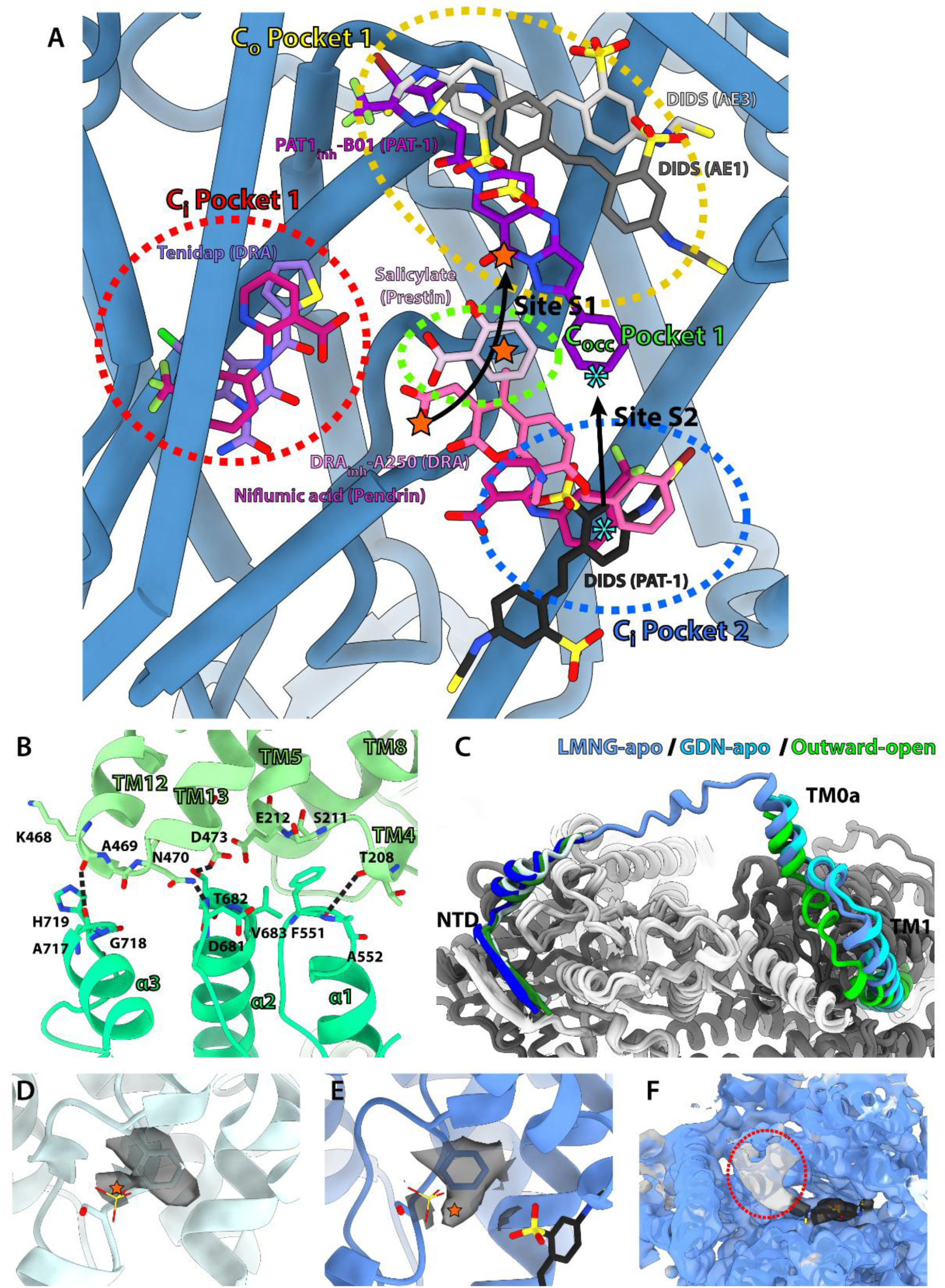
Further features of PAT-1 structures. **A)** Overview of inhibitor binding modes for SLC4 and SLC26 transporters. Two clusters of inhibitor pockets exist in inward-open anion transporters (Ci pockets 1 and 2). Ci pocket 2 also overlaps with Site S2. Occluded-state pocket (Cocc pocket 1) and outward-open inhibitor pocket (Co Pocket 1) overlap with Site S1. **B)** Interactions between TMD (light green) and STAS (green) near the dimer axis. **C)** Structure overview of NTD and NTD-TMD linker. Although TM1 of LMNG-apo structure is in the inward-open position (cyan), its TM0a is positioned like the outward-open conformation (green). **D)** Putative substrate density near F147 in Site S1 of GDN-apo PAT-1 structure. Sulphate from inward-open DRA (8X1T) is overlaid on the structure. Orange star denotes centre of the ion-like density. **E)** Putative substrate density in Site S1 of PAT-1:DIDS structure. **F)** Unmodelled large density on the cytoplasmic side of TMD of PAT-1:DIDS structure. Black – DIDS. Light grey density and red circle – unmodelled density.

**Supplementary Figure 5.**
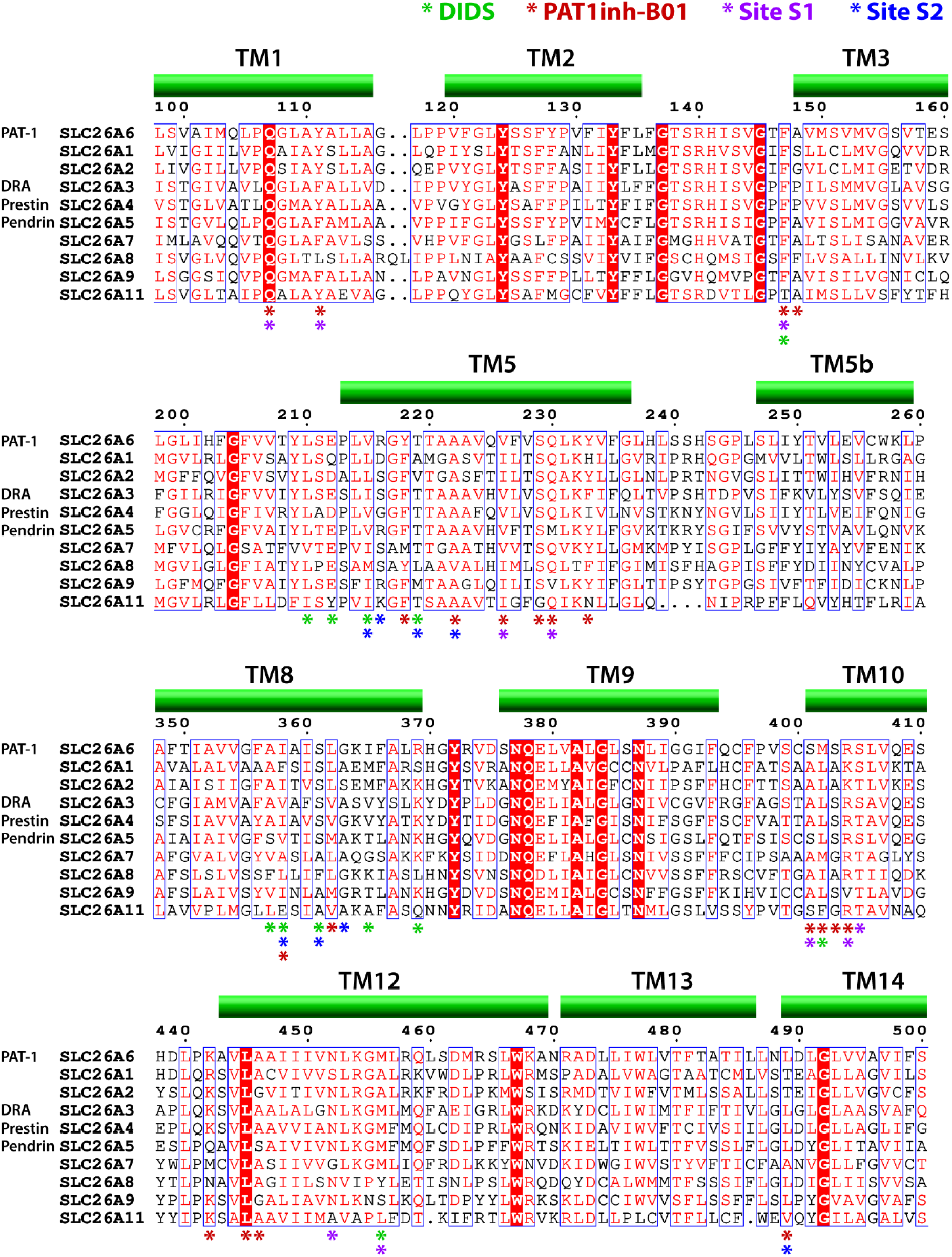
Sequence alignment of human SLC26 transporters. Asterisks denote PAT-1 residues comprising DIDS-binding site (green), PAT1_inh_-B01-binding site (red), substrate-binding Site S1 (purple), and Site S2 (blue).

**Supplementary Figure 6.**
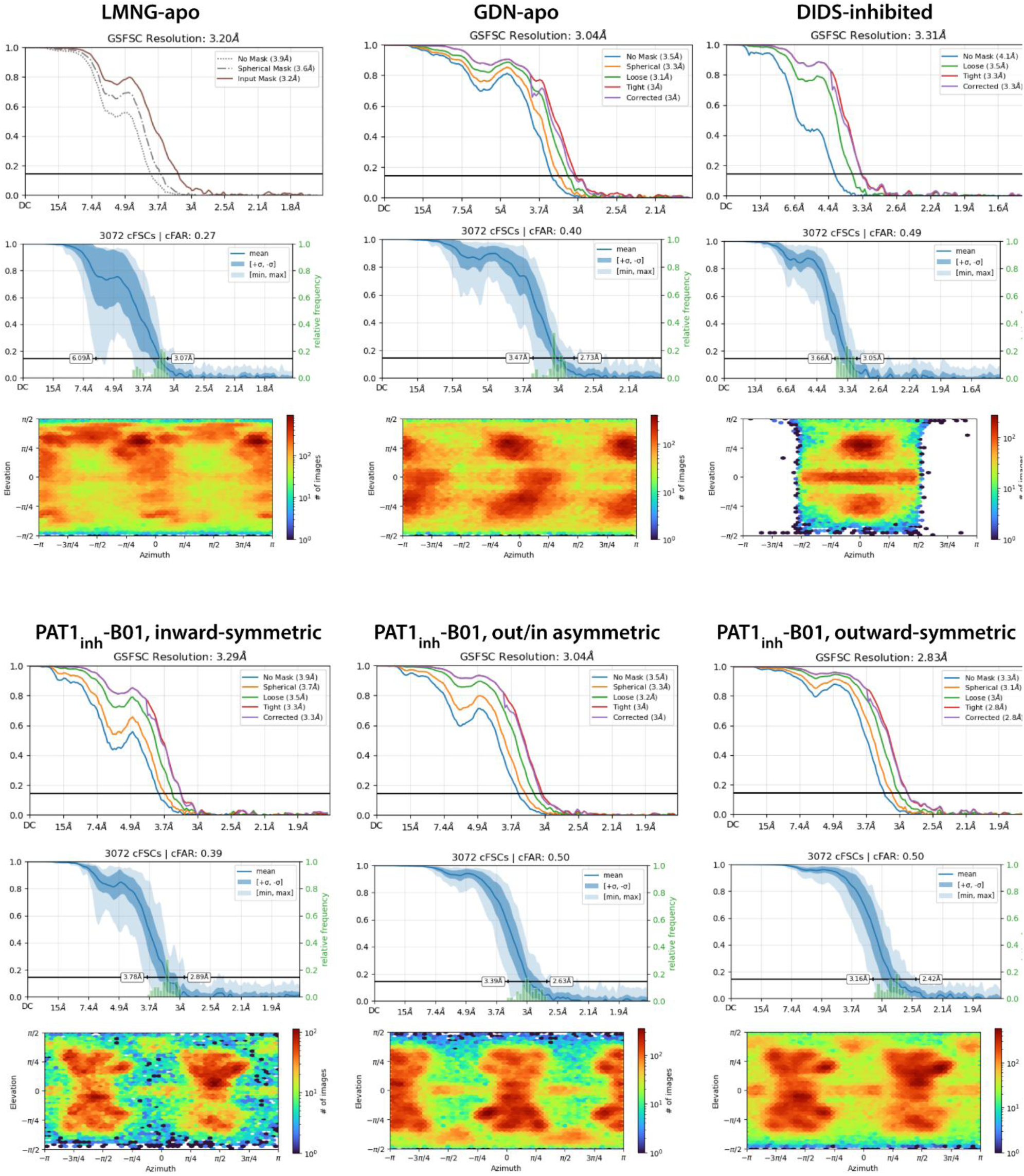
Fourier Shell Correlation (FSC) and cFSC charts, and orientation distribution plot for each PAT-1 cryo-EM dataset.

**Supplementary Figure 7.**
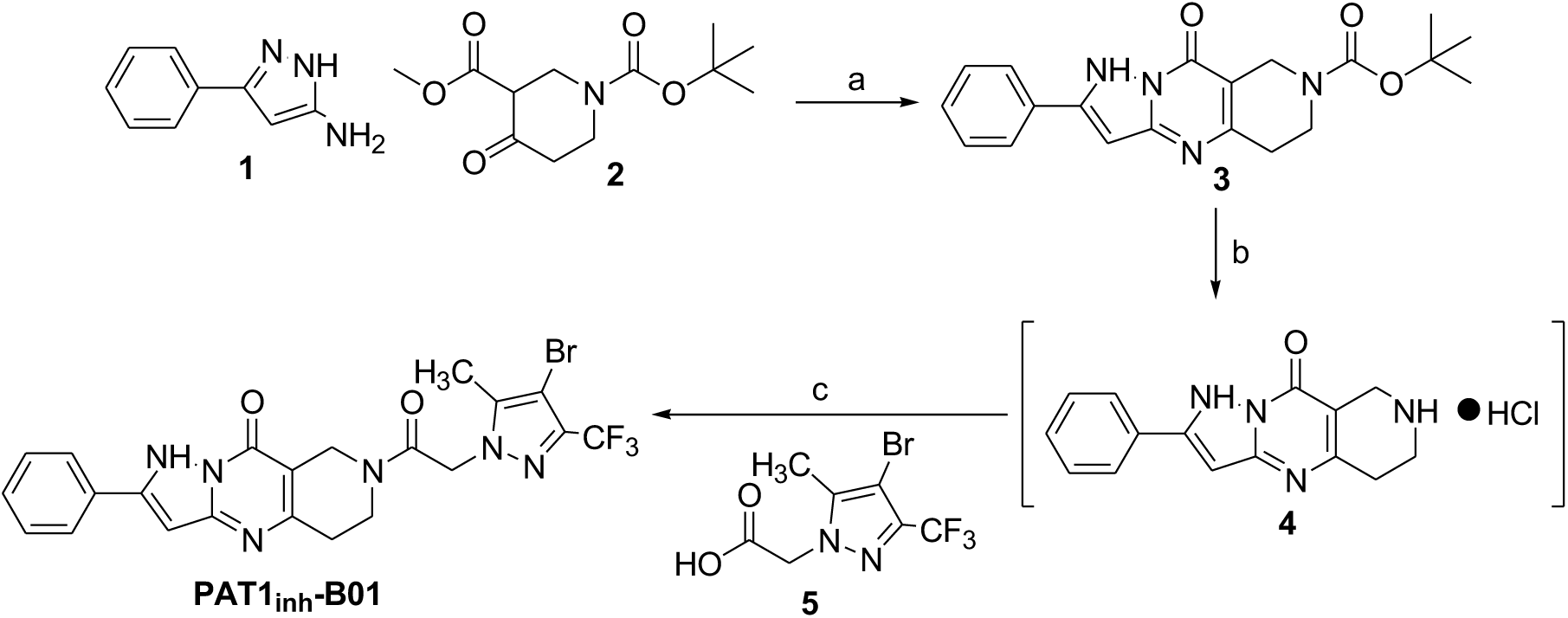
Synthesis of inhibitor PAT1inh-B01. Reagents and Conditions: (a) acetic acid, 100 ^0^C, 1.5h; (b) HCl (1M in EtOAc, 30 eq, RT, 18h); (c) EDCI, DCM, Et_3_N.

## Supplementary Methods

### Synthesis of PAT1_inh_-B01

Unless otherwise indicated, all reaction solvents were anhydrous and obtained as such from commercial sources. Chemical and solvents were commercially available and no further purification was required. Low resolution ESI-LCMS was carried out with an Agilent 1100 HPLC coupled to an Agilent 1956B MSD. High resolution ESI-MS was performed using an Agilent 1292 Infinity II UHPLC coupled to an Agilent 6546 LC/QTOF MSD instrument equipped with electrospray ionization. RP separation was achieved using an Agilent EclipsePlus C18 RRHD 1.8uM UHPLC column [2.1 x 50mm; Agilent part # 959757-902]. RP-HPLC runs typically employed gradients of two solvents: [A]= H2O (0.05% TFA) and [B] CH3CN (0.05% TFA); RP-LCMS used the same solvent system using the modifier formic acid (88% aq). The standard HPLC and LCMS gradients proceeded with [A:B]= 95:5 to [A:B]= 5:95 over 15 minutes. ^1^H and ^13^C NMR spectra were recorded on a Bruker 500 MHz instrument. ^1^H NMR chemical shifts are relative to TMS (δ = 0.00 ppm), CDCl_3_ (δ 7.26) or DMSO-*d_c_* (δ 2.5). ^13^C NMR chemical shifts are relative to CDCl_3_ (δ 77.2) or DMSO-*d_c_* (δ 39.5). Purity of assayed compounds was >95% based on HPLC-LCMS analysis at 254 nm, and absence of impurities was confirmed by examination at least one other wavelength (320 nm) as well as careful inspection of ^1^H and ^13^C-NMR spectra.

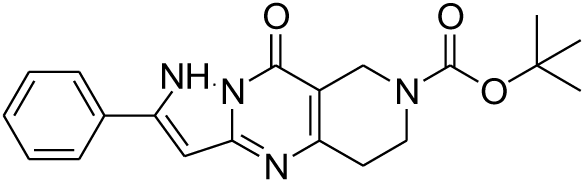

### G-Oxo-2-phenyl-1,5,6,G-tetrahydro-8H-1,4,7,Ga-tetraaza-cyclopenta[b]naphthalene-7-carboxylic acid *tert*-butyl ester (3)

Into a 20ml vial was added 1-*tert*-butyl 3-methyl 4-oxopiperidine-1,3-dicarboxylate (**2**) (1500mg, 5.83 mmol) and 3-amino-5-phenylpyrazole (**1**) (974mg, 6.12 mmol), which were dissolved in glacial AcOH (14.5ml). The reaction mixture was stirred and heated at 100 ^0^C for 1.5h, which generated substantial amounts of white precipitate. LCMS showed consumption of SM, and formation of product. The reaction mixture was added to an Erlenmeyer flask, washed over with Et_2_O, cooled in a freezer for 5min, and then vacuum filtered, to generate the product as a white crystalline solid. (1.931g, 90% yield). ^1^H NMR (500 MHz, DMSO-*d_c_*): δ 1.45 (s, 9H), 2.73 (t, 2H, *J* = 6 Hz), 3.63 (t, 2H, *J* = 6 Hz), 4.26 (s, 2H), 6.5 (s, 1H), 7.42 (t, 1H, *J* = 7 Hz), 7.48 (t, 2H, *J* = 7 Hz), 8.00 (d, 2H, *J* = 7 Hz). ^13^C NMR (125 MHz, DMSO-*d_c_*): δ 26.5, 28.5, 40.5, 79.9, 85.3, 126.7, 129.2, 129.4, 132.9, 142.7, 146.0, 153.7, 154.3, 155.6. ESI-LCMS (low resolution) m/z calculated for C_20_H_22_N_4_O_3_ [M+H] 367.1, found [M+Na] 367.1.

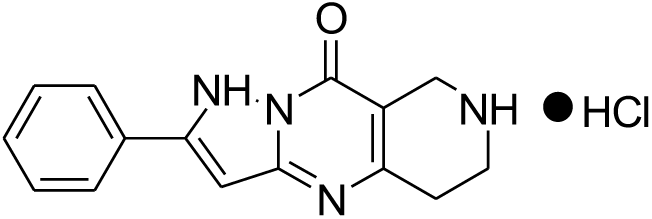

### 2-Phenyl-5,6,7,8-tetrahydro-1H-1,4,7,Ga-tetraaza-cyclopenta[b]naphthalen-G-one hydrochloride salt (4)

The starting material 9-oxo-2-phenyl-1,5,6,9-tetrahydro-8H-1,4,7,9a-tetraaza-cyclopenta[b]naphthalene-7-carboxylic acid tert-butyl ester (**3**) (200mg, 0.5 mmol) was dissolved in HCl (1M in EtOAc) (15ml, 30 eq) and stirred for 43h. LCMS showed consumption of SM and formation of deprotected product, which was added to an Erlenmeyer flask and treated with Et_2_O to assist in precipitation of the reaction mixture, which was vacuum filtered to generate the title product as a white crystalline solid (145mg, 88% yield). ^1^H NMR (500 MHz, DMSO-*d_c_*): δ 2.97 (t, 2H, J = 6 Hz), 3.40 (t, 2H, J = 6 Hz), 4.01 (s, 2H), 6.66 (s, 1H), 7.42 (t, 2H, J = 7 Hz), 7.49 (t, 2H, J = 7 Hz), 8.00 (d, 2H, J = 7 Hz), 9.37 (bs, 2H). ^13^C NMR (125 MHz, DMSO-*d_c_*): δ 23.4, 39.1, 85.9, 96.7, 126.7, 129.2, 132.6, 142.5, 144.6, 154.1, 155.2. ESI-LCMS (low resolution) m/z calculated for C_15_H_14_N_4_O [M+1] 267.1, found [M+1] 266.8.

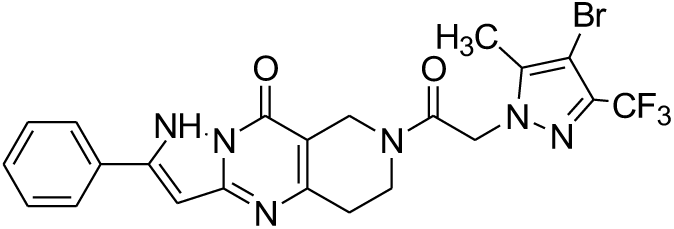

### 7-[2-(4-Bromo-5-methyl-3-trifluoromethyl-pyrazol-1-yl)-acetyl]-2-phenyl-5,6,7,8-tetrahydro-1H-1,4,7,Ga-tetraaza-cyclopenta[b]naphthalen-G-one (PATinh-B01)

The starting material 2-phenyl-5,6,7,8-tetrahydro-1H-1,4,7,9a-tetraaza-cyclopenta[b]naphthalen-9-one hydrochloride salt (**4**) (100mg, 0.33 mmol) was suspended in DCM (6.6ml) and treated with Et_3_N (0.048 ml, 0.34 mmol), followed by (4-bromo-5-methyl-3-trifluoromethyl-pyrazol-1-yl)-acetic acid (**5**) (151.7mg, 0.528 mmol) and EDCI-HCl (101.3 mg, 0.528 mmol). The reaction was stirred overnight at RT. The product was taken up into EtOAc (20ml), and washed with HCl (1M):brine [3:1 mixture] (2 x 10mL), dried over Na2SO4, and concentrated in vacuo to yield the crude product, contaminated with starting carboxylic acid (**5**). Removal of this contaminant was accomplished by trituration with Et_2_O, to yield the product as a white solid (29mg, 16% yield). ^1^H NMR (500 MHz, DMSO-*d_c_*): δ 2.23 (t, 3H), 2.76 (s, 1H), 2.95 (s, 1H), 3.79 (bs, 2H), 4.39 (s, 1H), 4.49 (s, 1H), 5.52 (s, 2H), 6.60 (s, 1H), 7.41-7.48 (m, 3H), 7.99-8.01 (m, 2H). ESI-LCMS (high resolution) m/z calculated for C_22_H_18_BrF_3_N_6_O_2_ [M+H] 535.0705, found [M+H] 535.0695.

